# Progression of Autoantibodies Anti-Gad and Anti-IA2 in Type 1A Diabetics Aged of 5 to 21 Years in Côte D’ivoire

**DOI:** 10.1101/551473

**Authors:** Aïssé Florence Judith Trébissou, Chiayé Claire Antoinette Yapo-Crezoit, Pascal Sibailly, Mamadou Sanogo, Amos Ankotché, Dinard Kouassi, Marie Thérèse Kouassi, Hatem Masmoudi, Adou Francis Yapo

**Affiliations:** Laboratory of Biochemistry-Pharmacodynamics, Faculty of Biosciences, Felix Houphouët Boigny University, Côte d’Ivoire, P.O BOX 582 Abidjan 22; National Public Health Institute, Abidjan, Côte d’Ivoire, P.O. BOX 47 Abidjan; Pole of Biology Immunity, Pasteur Institut of Côte d’Ivoire, P.O. BOX 490 Abidjan 01; Diabetes Clinic, University Hospital Center (U.H.C) of Treichville, Côte d’Ivoire; Diabetology and Endocrinology Department, University Hospital Center (U.H.C) of Yopougon; Laboratory of Immunology, Hospital Habib Bourguiba of Sfax, 3029Sfax, Tunisia

**Keywords:** diabetes autoantibodies, type 1A diabetes, Côte d’Ivoire

## Abstract

**Background:** Diabetes autoantibodies are indispensable markers of diabetes classification.

**Objective:** to research autoantibodies anti-GAD and anti-IA2 in type 1A diabetics (T1D) aged 5 to 21 years, and to follow the progression of these autoantibodies in T1D patients, in Côte d’Ivoire.

**Methods:** The study population composed of 28 T1D patients, aged 5 to 21 years. T1D were followed up in two diabetes care centers in Abidjan district, Endocrinology departments of U.H.C of Yopougon and Treichville. Anti-GAD and anti-IA2 autoantibodies were researched by ELISA.

**Results:** anti-GAD and anti-IA2 were present in T1D and their siblings. After 2 years of diabetes, the titer of the anti-GAD autoantibodies increased to the mean value of 677.10 ± 353.20 IU / ml. Then, a fall of the anti-GAD autoantibodies until the cancellation was observed from the 8th to the 9th with values of 117 IU / ml to 10.14 IU / ml. Anti-IA2 autoantibodies fall at 9th year of diabetes with a value of 55.10 IU / ml.

**Conclusion:** anti-GAD and anti-IA2 autoantibodies persist after 9 years of diabetes, causing total destruction over time of the pancreatic β-cell mass in patients from Côte d’Ivoire, leading them to the death.

## INTRODUCTION

Type 1A diabetes is an autoimmune disease, the final consequence of a slow and gradual process of ß-cell destruction of pancreatic islet Langerhans cells leading, in the absence of treatment, to ketoacidosis [1]. This destruction of the β cells, responsible for the production of insulin, begins with the initiation of the autoimmune reaction triggered by certain environmental factors and, after several years of evolution, leads to the clinical signs of the disease when the mass of ß cells become insufficient to regulate blood glucose [2]. Type 1A diabetes or juvenile insulin-dependent diabetes is a major public health problem due to the often early onset of the disease, increasing incidence in most populations, lack of curative treatment and vascular complications related to residual hyperglycemia despite insulin injections in treated patients [3]. It is a disease that affects more than 31 million people worldwide [4], and 138,000 people aged 19 to 79 died of type 1A diabetes in 2011 [5]. In Africa, 39100 children aged 0-14 years have type 1A diabetes in 2013 [5]. In Côte d’Ivoire, 0.4% of children and adolescents are type 1 diabetics (T1D) [6]. It is an incurable disease that is gaining momentum in Africa and particularly in Côte d’Ivoire. Today, the diagnosis of type 1A diabetes is biological (fasting glucose greater than 1.26 g / l) and requires the search for autoantibodies of diabetes as a confirmation marker for autoimmune process [7]. According to some authors, these autoantibodies diabetes disappear with the duration of diabetes [8]. In Côte d’Ivoire, diabetes autoantibodies screening tests are not performed leading to a lack of the diagnosis of type 1A diabetes and a high rate of morbidity and early mortality in this type of Ivorian population. The search for autoantibodies is essential in the classification of diabetics, some of whom are taken as type 2 diabetics when they are actually type 1A. It is therefore important to diagnose diabetes as early as possible in order to avoid the rapid evolution of the disease and its aggravation in the long term. The aim of this work was to research diabetes autoantibodies in T1D aged 5 to 21 years, and to follow the progression of these autoantibodies in T1D, in Côte d’Ivoire. This study was extracted from the Ph.D thesis written by Trébissou Aïssé Florence Judith.

## 1. Material and Methods

### 1.1 Material

#### Study population

The study population composed of 28 known T1D patients aged 5 to 21 years and followed up in the Endocrinology department of the University Hospital Center (U.H.C) of Yopougon and the Diabetes Clinic of U.H.C of Treichville, Abidjan, Côte d’Ivoire. The sex ratio was 0.86 (13 boys and 15 girls); and the mean age of T1D was 12.62 ± 2.75 years. This cross-sectional study started in January 2014 and ended in April 2016.

#### Selection criteria

##### 1. Inclusion criteria

The T1D selected were aged from 5 to 21 years and declared diabetics between 2007 and 2016. They had a complete medical folder of T1D.

##### 2. Non-inclusion criteria

HIV-positive patients undergoing antiretroviral therapy were excluded from the study because of the impact of ARVs that would lead to hyperglycemia.

#### Biological Material

The biological material composed essentially of serum. These sera were obtained after centrifugation at 3000 rpm for 3 minutes of venus blood collected on dry tubes in T1D and their siblings.

### 1.2 Methods

The project was approved by the ethics committee of Pasteur Institut of Côte d’Ivoire.

#### 1.2.1 Search of Anti-Glutamic Acid Decarboxylase Autoantibodies and anti-tyrosine Phosphatase IA2 Autoantibodies

The dosage of these two autoantibodies is done by ELISA method with commercial kits anti-GAD ELISA (IgG) and anti-IA2 ELISA (IgG) (Euroimmun^®^, Germany). The test is positive if the titer of anti-GAD and anti-IA2 autoantibodies is greater than or equal to 10 IU/ml [9].

### 1.3 Statistical analysis

Statistical analysis of the data was done with the GraphPad Prism.V5.01 software using the Student’s test. The difference between two variances was significant if p<0.05.

### 1.4 Results

Figure 1 shows the titer of anti-GAD autoantibodies according to the age of the disease in T1D. These results show that after one year of diabetes, the titer of anti-GAD autoantibodies was 63.30 ± 37.70 IU/ml. After 2 years of diabetes, the mean value of the anti-GAD autoantibodies raised to 677.10 ± 353.20 IU / ml (Figure 3). After 5 to 6 years of diabetes, the results showed a fall in anti-GAD autoantibodies from 587.60 ± 305.80 IU / ml to 247.40 ± 210.70 IU / ml. Then a rebound of anti-GAD autoantibodies was observed at the 7th year of diabetes with a value of 956.50 ± 171.50 IU / ml. Then, a fall of the anti-GAD autoantibodies until the cancellation was observed from the 8th to the 9th year with values of 117 IU / ml to 10.14 IU / ml (Figure 1).

**Figure 1:**
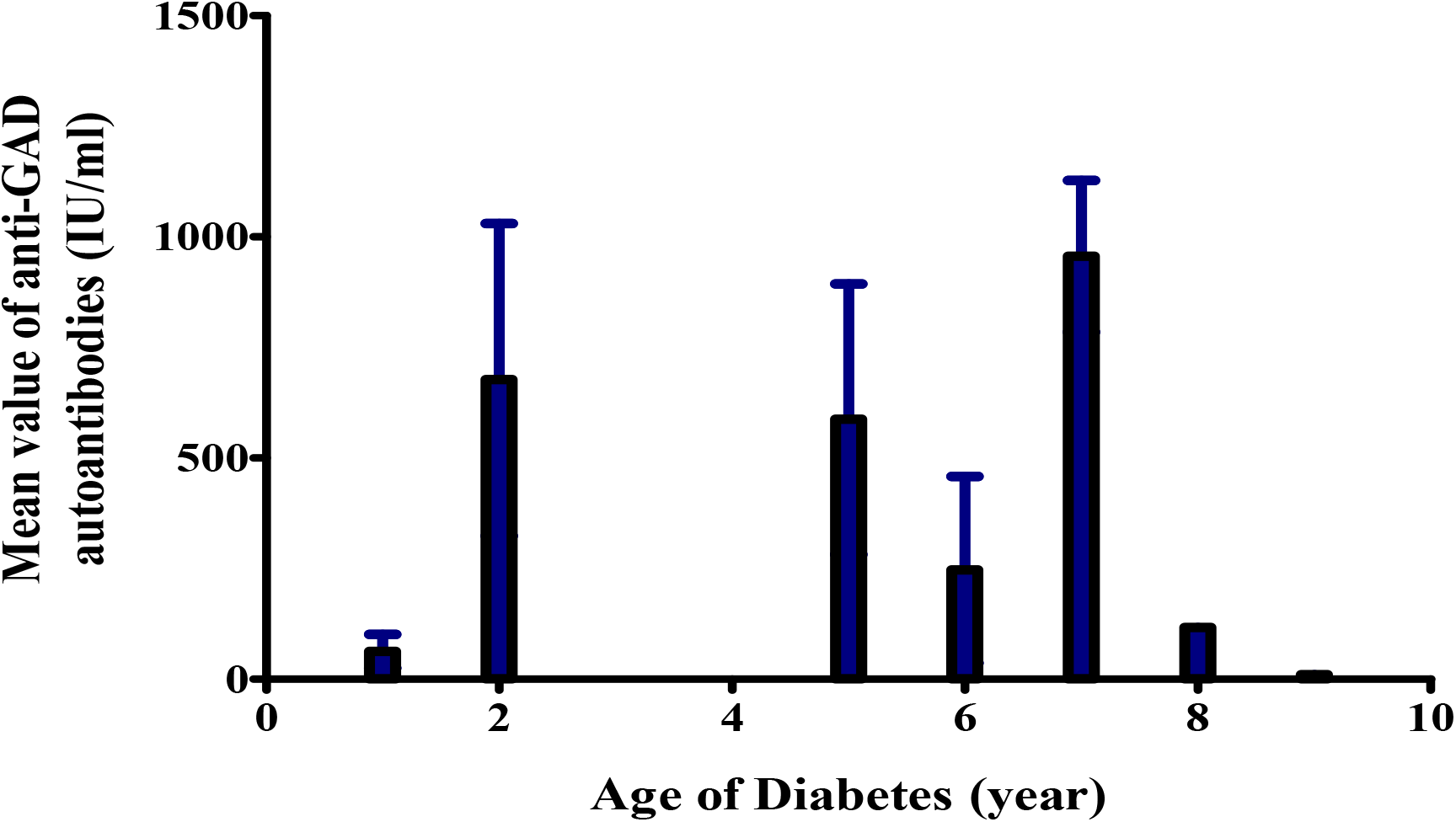
Mean value of Anti-GAD autoantibodies according to age of diabetes

Figure 2 shows the titer of anti-IA2 autoantibodies according to the age of diabetes in T1D. These results show that after one year of diabetes, the titer of the anti-IA2 autoantibodies was at its maximum value of 321 IU / ml. Then, a fall of the anti-IA2 autoantibodies from the 2nd year until the 7th year was observed, with values varying from 248,60 ± 95,40 IU / ml, 89,80 ± 25,78 IU / ml, 49.95 ± 39.05 IU / ml at 25.73 IU / ml (2nd, 5th, 6th and 7th year of diabetes). Subsequently, a slight increase in anti-IA2 autoantibodies was observed in the 9th year of diabetes with a value of 55.10 IU / ml (Figure 2).

**Figure 2:**
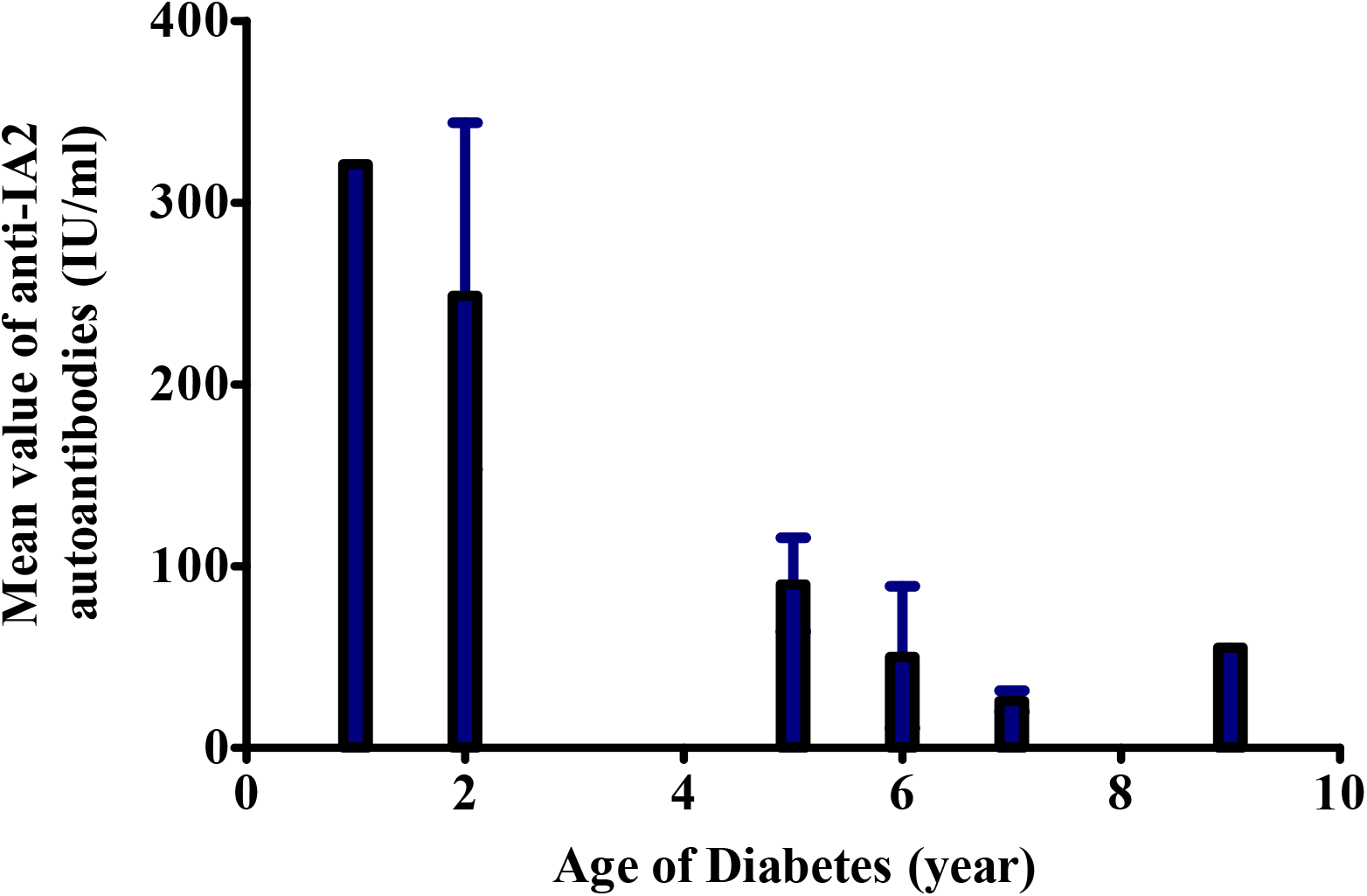
Mean value of Anti-IA2 auto-antibodies according to age of diabetes

## Discussion

Type 1A diabetes is an autoimmune disease due to the destruction of pancreatic islet β-cells by T cells. So, by a rupture of the peripheral tolerance of the immune system with respect to β cells-self-antigens, a phenomenon of “molecular mimicry” occurs between a sequence of a viral protein and that of an auto-antigen of the β-cell of Langerhans islets of the endocrine pancreas. Another mechanism involved is the « bystander effect », ie infection of the β-cells by a virus with a particular tropism for the latter, followed by their lysis and the release of self-antigens susceptible to activate self-reactive T cells for these autoantigens [10]. Thereby, T1D with young diabetes (1 to 2 years) who have not developed at least one diabetes autoantibody would not be T1D. So, as some authors have shown, diabetes autoantibodies disappear with duration of diabetes, hence the disappearance in time of diabetes auto-antibodies in T1D in Côte d’Ivoire. This would be due to the destruction of β-cells [8]. It appears important to introduce in diabetes care in Côte d’Ivoire the search for autoantibodies of diabetes, for a better classification and diabetes care.

The results so showed that the mean value of anti-GAD autoantibodies after 2 years of diabetes was maximal in T1D (677.10 ± 353.20 IU / ml). Then, from the 5th to the 6th year, it falled to reach 247.40 ± 210.70 IU / ml after 6 years of diabetes, followed by a brutal fall between the 8th and 9th year of diabetes (117 IU/ml at 10.14 IU/ml). With regard to anti-IA2 autoantibodies, the results showed a maximum value of 310 IU/ml in the first year of diabetes, followed by a brutal fall from the 2nd year, 5th, 6th until the 7th year (248.60 ± 95.40 IU/ml, 89.80 ± 25.78 IU/ml, 49.95 ± 39.05 IU/ml and 25.73 IU/ml). These results show that β-cells of Langerhans islets of the endocrine pancreas of these T1D know undergo a fast destruction, which can lead to the death of these. The rebound of anti-GAD autoantibodies observed at the 7th year of diabetes and at the 9th year for anti-IA2 is thought to be due to the persistence of the humoral immune response against the islet cells of the Langherans of the pancreas. Indeed, according to Wilmot-Roussel *et al*., 2010 [8] anti-GAD and anti-IA2 autoantibodies persit during the type 1A diabetes. The body of T1D has secreted more anti-GAD and anti-IA2 autoantibodies to destroy the remaining β cell mass.

## Conclusion

In conclusion, this study shows the important role that anti-GAD and anti-IA2 autoantibodies play in diabetes care. Indeed, anti-GAD and anti-IA2 autoantibodies persist after 9 years of diabetes. Anti-IA2 auto-antibodies would be more persistent than anti-GAD in T1D in Côte d’Ivoire. It appears essential to introduce the research of auto-antibodies anti-GAD and anti-IA2 in diabetes care in Côte d’Ivoire, to allow better monitoring of diabetics.

## ACKNOWLEDGEMENTS

We thank the Laboratory of Immunology, Hospital Habib Bourguiba of Sfax, 3029Sfax, Tunisia, for the financially supported and autoantibodies assay.

## AUTHOR CONTRIBUTIONS

**Conceptualization**: Aïssé Florence Judith Trébissou, Adou Francis Yapo, Chiayé Claire Antoinette Yapo-Crezoit.

**Data curation**: Aïssé Florence Judith Trébissou

**Formal analysis**: Aïssé Florence Judith Trébissou, Adou Francis Yapo

**Funding acquisition**: Hatem Masmoudi

**Investigation**: Aïssé Florence Judith Trébissou, Sibailly Pascal, Sanogo Mamadou, Ankotché Amos

**Methodology**: Aïssé Florence Judith Trébissou

**Supervision**: Kouassi Dinard, Kouassi Marie Thérèse

**Writing – original draft**: Aïssé Florence Judith Trébissou

**Writing – review & editing**: Aïssé Florence Judith Trébissou, Adou Francis Yapo

